# Uncovering asparagine utilization pathway, a critical requirement for mycobacterial pH-driven adaptation at the host pathogen crossroads

**DOI:** 10.1101/2025.06.09.658563

**Authors:** Bhanwar Bamniya, Kajal, Khushboo Mehta, Dibyendu Sarkar

## Abstract

*M. tuberculosis*, an intracellular pathogen, thrives in a specialized membrane-bound vacuole, the phagosome with acidic pH and limited access to nutrients. To survive and replicate within human host, *M. tuberculosis* must adapt and fulfil its nutritional requirements. To understand how the bacillus captures nutrients from its host, we studied mycobacterial acquisition of host-derived amino acids, the preferred nitrogen sources. We discovered that PhoP, a key determinant of mycobacterial adaptation to phagosomal acidification, controls expression of AnsP1 and AnsP2 to facilitate acquisition of host aspartate and asparagine, respectively. Thus, macrophage-infected WT-H37Rv showed a significantly higher level of intra-bacterial Asn compared to the *phoP*-KO mutant and a complemented mutant could restore Asn level to that of WT-H37Rv. Under acidic conditions, elevated DNA binding of PhoP within the promoters lead to direct activation of *ansP1* and *ansP2*. Consistently, *phoP*-KO is unable to utilize Asn under acidic condition, and over-expression of *ansP1* or *ansP2* restored intracellular growth defect of the mutant. These findings uncover the regulatory network allowing utilization of organic nitrogen sources by the pathogen during infection.

**Author summary:** Human Tuberculosis (TB), caused by *Mycobacterium tuberculosis*, remains a global public health burden claiming ∼1.25 million deaths in 2023. To investigate how the bacillus captures nitrogen sources from its host, we studied acquisition of host amino acids aspartate (Asp) and asparagine (Asn) by the tubercle bacilli. We show that PhoP, a key determinant of mycobacterial adaptation to phagosomal acidification, controls expression of Asp and Asn transporters to facilitate acquisition of host amino acids by the pathogen. A mutant lacking *phoP* is unable to utilize Asn under acidic conditions, whereas over-expression of *ansP1* and/or *ansP2* restored growth defect of the *phoP*-KO *in vitro* and in macrophages. Together, these findings uncover the transcriptional landscape allowing utilization of organic nitrogen sources by the tubercle bacilli during infection.

## Introduction

Human tuberculosis is caused by the bacillus *Mycobacterium tuberculosis*, which thrives inside the host macrophages and evades eradication by the human immune system. Like other bacterial diseases, tuberculosis continues to become increasingly drug resistant with strains resistant to two frontline drugs (MDR), and an additional resistance to an injectable antibiotic and a quinolone (XDR), claiming lives of 500 thousand patients a year (WHO, 2017). Thus, the consensus opinion is that a better understanding of the interaction of the pathogen and its human host, particularly uncovering microbial mechanisms involved in acquisition and metabolism of host-derived nutrients by the pathogen during its complex life cycle may help identify new targets for novel antimicrobials [1–3]. As an intracellular pathogen, *M. tuberculosis* resides and multiplies inside a macrophage phagosome, which fuse poorly with host lysosomes [4–7]. However, a hallmark of virulence strategy by the pathogen is its unique ability to block phagosome maturation and avoid lysosomal degradation. While the molecular mechanism could be the resultant of a complex set of multiple events, our failure to identify transformative antitubercular therapy is largely attributable to our inadequate understanding of bacterial acquisition of host-derived nutrients by the pathogen.

The mycobacterial phagosome represents an environment comprising different niches with acidic pH, hypoxia, scarcity of nutrients and presence of reactive oxygen and nitrogen species [8, 9]. A number of high throughput studies showing induction of acid inducible genes and genes involved in utilization of alternative carbon sources, for example, host derived fatty acids and cholesterol [10–12], suggest that such an environment with multiple environmental cues translate into a significant remodeling of mycobacterial transcriptional landscape soon after phagocytosis. Previously, we had shown that *M. tuberculosis* has evolved mechanisms integrating hypoxia and metabolism of nitrogen, the alternate electron acceptor under limiting oxygen availability [13]. While this pathway promotes intracellular survival and growth of the bacilli in the phagosomal hypoxic environment, nitrogen remains an essential component of amino acids, nucleotides and many biomolecules like organic co-factors. Although most of the previous studies examining growth and metabolism utilized ammonium (NH ^+^) as the physiologically relevant nitrogen sources, there has been compelling evidence on the critical requirement of amino acids during *M. tuberculosis* infection [14, 15]. To identify host-relevant nitrogen metabolism in *M. tuberculosis*, two previous reports demonstrated that host-derived amino acids L-aspartate (Asp) and L-asparagine are critical nitrogen sources during infection [16, 17]. An elegant extension of these studies more recently has explored intricate details of the nitrogen metabolic network of *M. tuberculosis* [18]. These results demonstrate (a) a more efficient nitrogen utilization from proteinogenic amino acids compared to ammonium, (b) comparable rate of utilization of amino acids as nitrogen sources and (c) lack of stringent homeostatic control mechanism of nitrogen utilization using amino acids. What remains unknown is the transcriptional regulatory network that couples effective utilization of Asn as the most preferred nitrogen source under acidic pH conditions of growth.

*M. tuberculosis* lacks homologues of nearly all known bacterial transcription factors involved in nitrogen metabolism [19]. Thus, transcription regulation of nitrogen metabolism in *M. tuberculosis* is poorly understood. Given the fact that Asn, as one of the most preferred amino acids [19] provides mycobacterial resistance to low pH conditions of growth [20] in one hand, and mycobacterial adaptation to the phagosomal low pH conditions require functioning of the *phoPR* system on the other hand, we sought to investigate whether *phoP* locus is linked to Asn utilization of mycobacteria. We explored utilization of nitrogen by *M. tuberculosis* metabolic network with amino acids as sole source of nitrogen, employing bacterial physiology, transcriptional regulation and intracellular survival experiments. Our results demonstrate that virulence regulator PhoP functions as a DNA binding transcription factor to impact expression of Asp transporter, AnsP1 and Asn transporter, AnsP2, respectively. Along the line, acidic pH-dependent elevated DNA binding by PhoP within the promoters of Asn/Asp transporters contributes to restructure transcriptional landscape, impacting mycobacterial tolerance to acidic pH conditions and intracellular survival within the host. In keeping with these results, mycobacterial growth inside phagosomes is supported by release of ammonia and consequent pH buffering. These results uncover a novel mechanism of how phagosomal adaptation of mycobacteria [21, 22] integrates nitrogen metabolism and acidic conditions of growth to promote intracellular survival of the pathogen.

## RESULTS

### Asn is an effective nitrogen source of *M. tuberculosis* growth under acidic conditions

A recent study elegantly demonstrated that *M. tuberculosis* has a very flexible metabolic network with Asn and Gln as the most preferred nitrogen sources [18]. To examine the effect of nitrogen source on mycobacterial growth, we attempted to characterize *in vitro* growth phenotypes of the *M. tuberculosis* wild-type (WT-H37Rv) using synthetic Middlebrook 7H9 (m7H9) liquid medium containing either Asn or Gln as the sole source of nitrogen (Figs. 1A-B). Growth kinetics of WT-H37Rv was almost identical with either of the nitrogen sources both under normal (pH 6.6) and acidic conditions (pH 5.5). Enumeration of viable CFU counts showed that WT-H37Rv grows at comparable efficiency both under normal and acidic conditions in m7H9 supplemented with Asn as the sole nitrogen source (Fig. 1C). However, utilization of Gln as the sole source of nitrogen under acidic conditions was less effective for WT-H37Rv relative to normal conditions of growth. As controls, growth kinetics of WT-H37Rv in m7H9 supplemented with ammonium salt or Asp showed significantly reduced growth of the bacteria under acidic conditions relative to normal conditions (Figs. S1A and S1B, respectively). Consistent with growth kinetics, viable CFU counts displayed similar results with a significantly reduced growth of WT-H37Rv under acidic pH relative to normal conditions (Figs. S1C and S1D, respectively).

**Figure 1:**
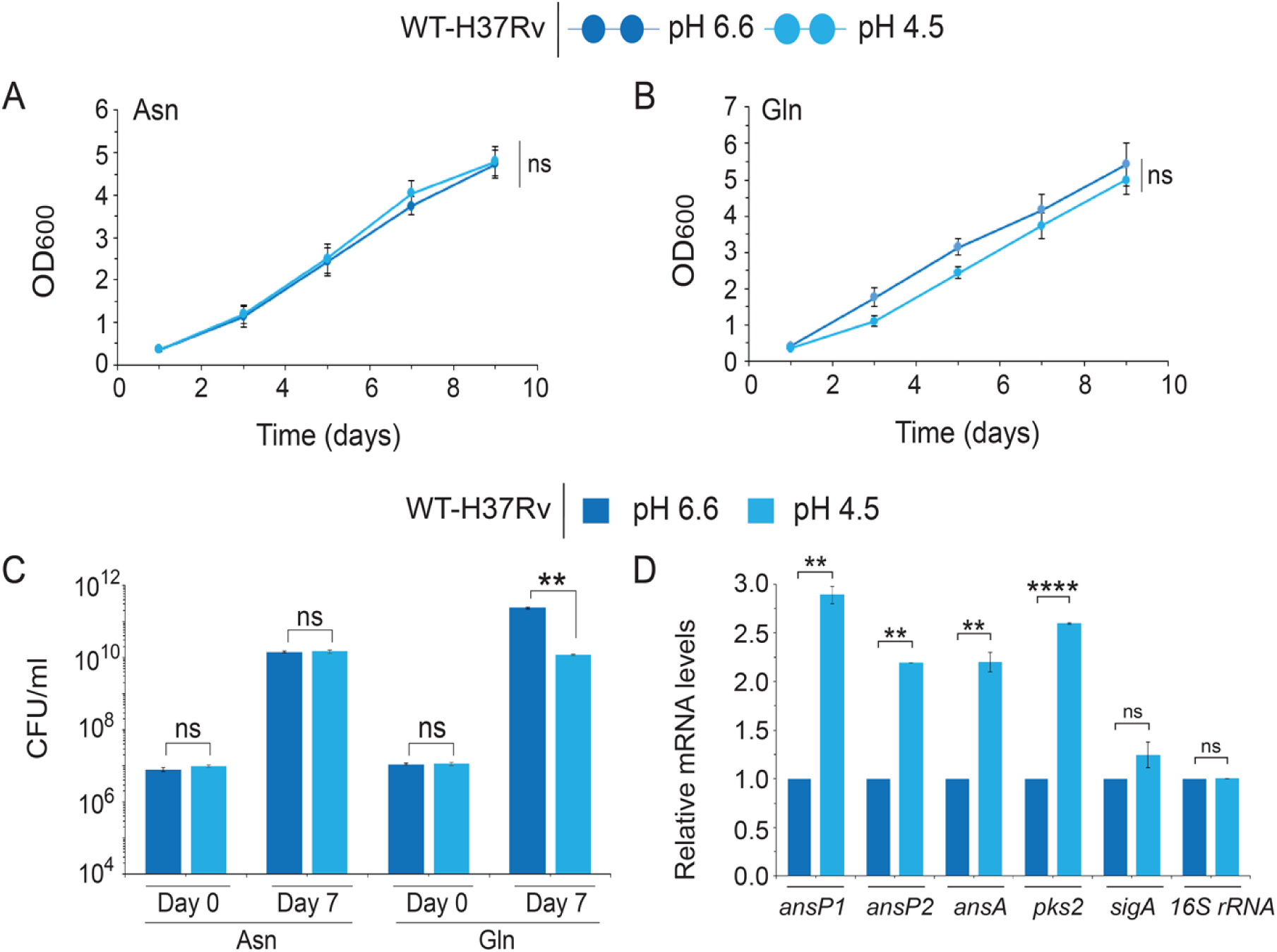
Effective utilization of Asn by *M. tuberculosis* under acidic conditions of growth. To examine mycobacterial utilization of amino acids as the nitrogen source, WT-H37Rv was grown *in vitro* in synthetic m7H9 medium supplemented with (A) Asn or (B) Gln as the sole nitrogen source under pH 6.6 (normal conditions) and pH 4.5 (acidic conditions), respectively. The growth experiments were performed in biological triplicates, each with technical repeats. Panel (C) enumerates CFU values for both normal and acidic conditions at day 0 and day 7, respectively (** P<0.01; ns, non-significant difference). (D) To investigate expression of genes related to Asn metabolism, expression of representative genes was compared in WT-H37Rv grown under normal (pH 6.6) and acidic pH conditions (pH 4.5). The data show plots of average values from biological triplicates, each performed with technical repeats (****P≤0.0001; **P≤0.01; *P≤0.05; ns, non-significant difference).

Together, these results suggest that Asn is an effective nitrogen source, supporting mycobacterial growth under acidic pH.

Having shown effective Asn utilization by mycobacteria grown under acidic pH, we undertook a transcriptomic approach to investigate expression of genes related to Asn utilization. To this end, we grew WT-H37Rv under normal and acidic pH conditions of growth and assessed relative abundance of mRNAs related to Asn utilization (Fig. 1D). We observed low pH-inducible expression of aspartate transporter *ansP1*, asparagine transporter *ansP2*, and asparaginase *ansA*, respectively. *pks2*, encoding low-pH inducible polyketide synthase was used as a positive control, whereas *sigA* and *16S rRNA* genes served as endogenous controls. Notably, low pH-inducible expression of mycobacterial *ansP1*, *ansP2*, and *ansA* is consistent with previous studies suggesting importance of L-Asparatate (Asp) and L-asparagine (Asn) as critically important nitrogen sources [16, 17] during infection and the role of Asn in mycobacterial resistance to low pH conditions [20]. These observations are consistent with previously reported results suggesting Asn as the most effective nitrogen source of mycobacteria [18] and Asn-dependent mycobacterial resistance to acidic stress [20].

### Asn utilization of *M. tuberculosis* requires the *phoP* locus

PhoP has been implicated in mycobacterial adaptation to low pH conditions inside the macrophage phagosome and is significantly activated under acidic pH both *in vitro* and *ex vivo* [21–24]. Since Asn was effectively utilized by mycobacteria under normal and acidic conditions of growth with comparably efficiency, we next explored role of the *phoP* locus in mycobacterial Asn utilization. We adopted a previously-reported CRISPR interference-based approach [25] to generate a *phoP* knock-down (*phoP*-KD) construct as described earlier [26]. To examine *in vitro* growth phenotype, *phoP*-KD mutant was grown under normal conditions in synthetic m7H9 containing Asn as the sole nitrogen source (Fig. 2A). Although *phoP*-KD mutant showed a significantly reduced growth relative to WT-H37Rv, under identical conditions a control strain (*prrB-*KD) showed a similar growth profile as that of WT-H37Rv. However, both knock-down mutants grew as effectively as the WT-bacilli with Gln (Fig. 2B) or ammonium (Fig. S2A) as the sole nitrogen source. It should be noted that *prrB*-KD strain showed a reduced *prrB* expression without any significant change in *phoP* expression level (Fig. S2B).

**Figure 2:**
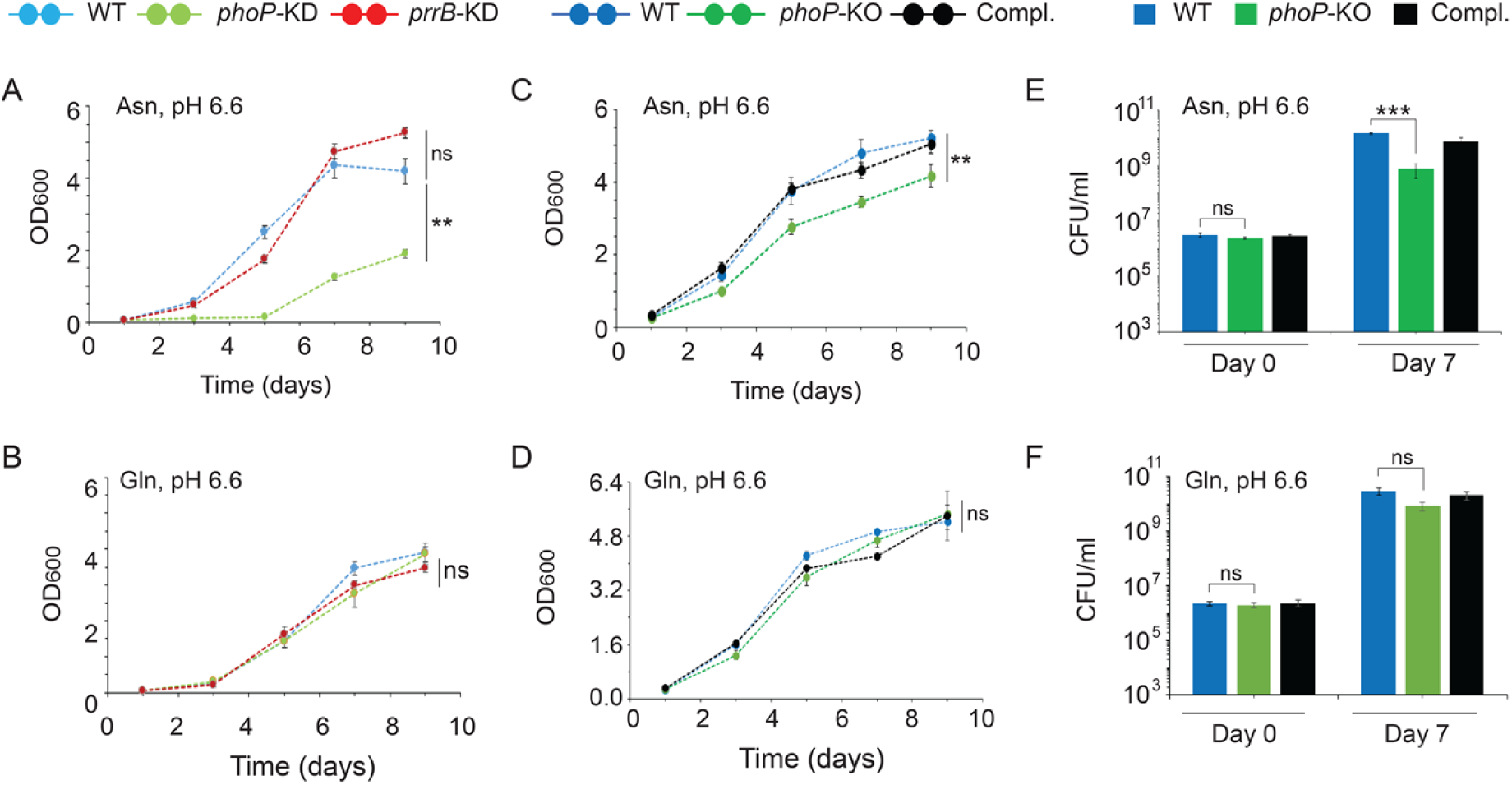
Asn utilization by *M. tuberculosis* requires the *phoP* locus. To determine the role of mycobacterial *phoP* locus, growth experiments of WT-H37Rv and indicated knock-down mutants were performed in synthetic m7H9 medium supplemented with (A) Asn, or (B) Gln as the sole nitrogen source, under normal conditions. (C-D) Identical growth experiments were also carried out with WT, *phoP*-KO and the complemented mutant in synthetic m7H9 using (C) Asn or (D) Gln as the sole nitrogen source. (E-F) Enumeration of CFU of WT, *phoP*-KO and the complemented mutant strain (Compl.) under normal conditions at day 0 and day 7, respectively. These experiments were performed in biological triplicates, each with two technical repeats, (***P<0.001; ns, non-significant difference).

To further probe the role of *phoP* locus on Asn utilization, we next grew WT-H37Rv, *phoP*-KO and complemented *phoP*-KO mutant (Compl.) in synthetic m7H9 supplemented with Asn or Gln as sole source of nitrogen (Fig. 2C-E). Our results demonstrate that *phoP*-KO is significantly growth defective in synthetic m7H9 media containing Asn as the sole nitrogen source (Fig. 2C). However, under identical conditions, complemented mutant expressing a copy of the *phoP* gene showed a largely similar growth as that of WT-H37Rv. In sharp contrast, WT bacteria, *phoP*-KO mutant and the complemented mutant strain displayed a comparable growth profile in m7H9 supplemented with either Gln (Fig. 2D) or ammonium (Fig. S2C) as the sole source of nitrogen. These results further suggest that Asn utilization of mycobacteria requires the *phoP* locus. Notably, CFU enumeration of mycobacterial strains grown in synthetic m7H9 supplemented with Asn or Gln as the sole nitrogen source (Figs. 2E and 2F, respectively) displayed consistent results. Together, these results uncover a striking effect of *phoP* locus on mycobacterial utilization of Asn, but not Gln.

### pH homeostasis is linked to mycobacterial Asn utilization via the *phoP* locus

Previous studies implicate a major role of the response regulator PhoP in phagosomal pH-dependent adaptation of mycobacteria[21, 22, 24]. While low pH remains the activating signal of *phoR-phoP* system [23], results reported here suggest that Asn utilization of mycobacteria requires the *phoP* locus. To probe whether intra-bacterial pH homeostasis is linked to mycobacterial Asn utilization, we have transformed WT-H37Rv and the *phoP*-KO strain with a GFP-based pH sensitive probe [27], and grew the strains both under acidic (pH 5.5) and normal (pH 6.6) conditions using Asn as the sole nitrogen source. Based on a standard curve plotting intra-bacterial pH as a function of fluorescence intensities from bacterial cells grown under varying pH conditions [27], we could accurately determine intra-bacterial pH. WT bacteria, under acidic conditions of growth, could maintain an intra-bacterial pH of ∼6.60 (Fig. 3A). However, under identical conditions, *phoP*-KO failed to maintain intra-bacterial pH. As a control, we did not observe a significant difference in intra-bacterial pH between WT-H37Rv and the *phoP*-KO mutant grown in the presence of ammonium as the nitrogen source. These results are consistent with Asn acquisition-dependent intra-bacterial pH homeostasis via mycobacterial *phoP* locus.

**Figure 3:**
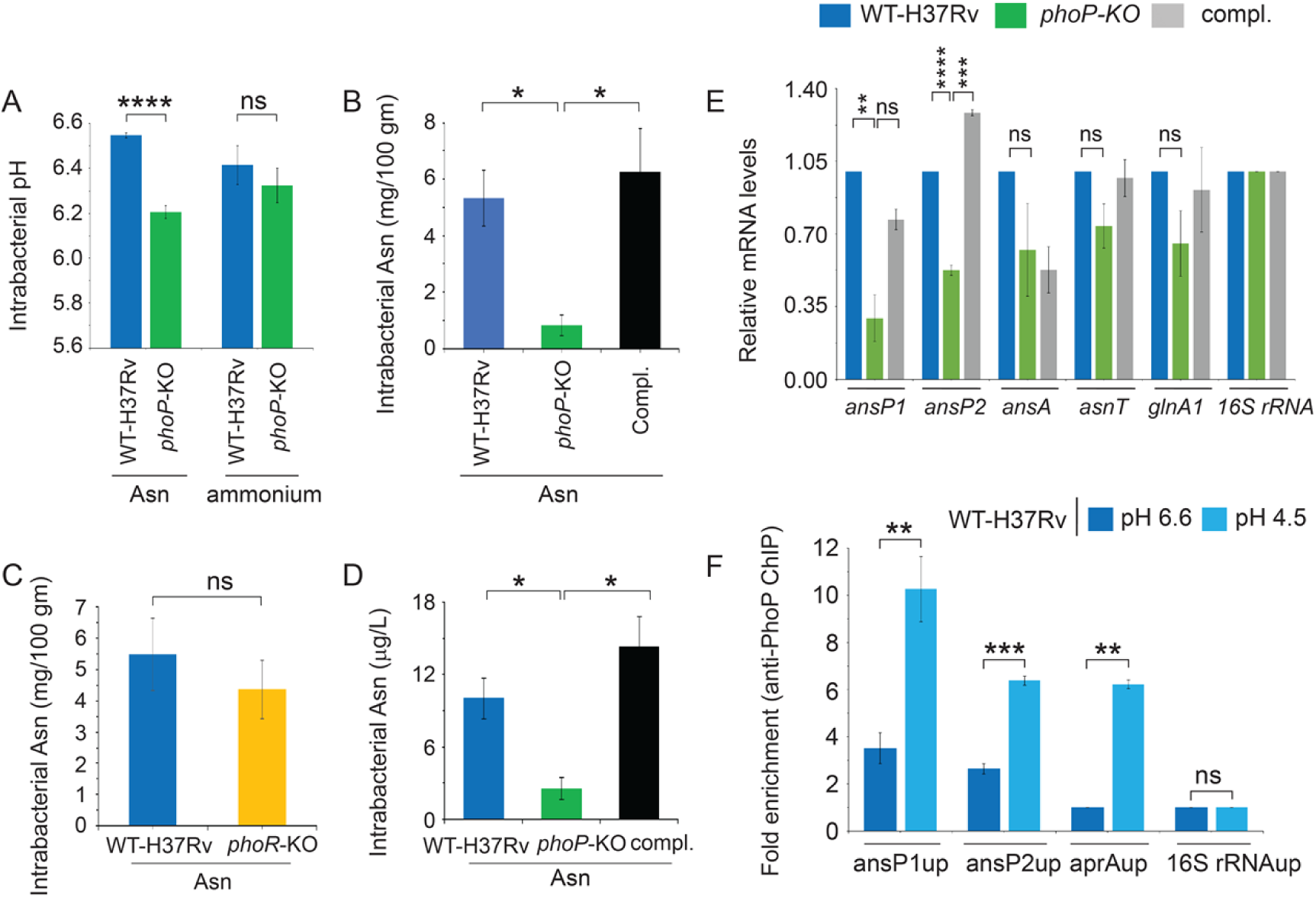
Asn utilization and pH homeostasis are linked by the *M. tuberculosis phoP* locus. (A-B) WT-H37Rv and *phoP*-KO mutant were grown under acidic pH conditions (pH 4.5) using synthetic m7H9 medium supplemented with indicated nitrogen source, and intra-bacterial (A) pH, and (B) Asn measurements were performed as described in the methods. As a control, panel (C) compares intra-bacterial Asn levels of WT-H37Rv and the *phoR*-KO mutant, grown under identical conditions. (D) To investigate Asn acquisition from macrophages *in vivo*, THP-1 macrophages were infected with WT-H37Rv, *phoP*-KO mutant, and the complemented mutant strain, and 48-hour post-infection bacterial strains isolated from infected macrophages, and intra-bacterial Asn levels measured as described in the methods. The values shown here represent the average of biological triplicates each with technical repeats (****P< 0.0001; *P≤0.05; ns, non-significant difference). (E) Expression of low pH inducible mycobacterial genes related to Asn metabolism were compared in WT-H37Rv, *phoP*-KO, and the complemented mutant grown under acidic conditions (pH 4.5) and in the presence of Asn as the sole nitrogen source using RT-PCR, as described in the methods. (F) To examine *in vivo* recruitment of PhoP within the relevant promoters, ChIP was carried out using anti-PhoP antibody followed by qPCR using IP samples from WT-H37Rv grown under normal (pH 6.6) and acidic conditions (pH 4.5). Each data point was compared with the corresponding IP sample without adding antibody (mock sample) to determine fold enrichment. For both RT-qPCR and ChIP-qPCR studies the data represent average of biological triplicates, each performed with technical repeats (****P≤0.0001; ***P≤0.001; **P≤0.01; *P≤0.05; ns, non-significant difference).

In an identical experimental set up, we next compared intra-bacterial Asn level of WT-H37Rv and *phoP*-KO mutant (Fig. 3B). Our results demonstrate a significantly lower Asn level of the *phoP*-KO mutant relative to WT bacteria; however, stable expression of a copy of *phoP* gene in the mutant restored Asn level to the WT bacilli. These results suggest that intra-bacterial pH homeostasis of mycobacteria is linked to Asn acquisition via the *phoP* locus. These observations suggesting specific role of *phoP* locus received further support with no significant difference in intra-bacterial Asn levels when compared between WT-H37Rv and *phoR*-KO mutant, grown under identical conditions in the presence of Asn as the sole nitrogen source (Fig. 3C). In agreement with these results, under identical conditions we observed a significantly lower level of intra-bacterial ammonia in *phoP*-KO mutant relative to the WT bacilli and the complemented mutant (Compl.) (Fig. S3A).

To investigate mycobacterial Asn acquisition from macrophages *in vivo*, we infected THP-1 macrophages with WT-H37Rv and *phoP*-KO mutant of *M. tuberculosis*. 48-hour post infection, bacteria were isolated from infected macrophages, macrophage debris removed by extensive washing and intra-mycobacterial Asn levels measured from comparable counts of bacterial cells. Under identical conditions, macrophage-derived WT-H37Rv showed a significantly higher level of intra-bacterial Asn compared to the *phoP*-KO mutant (Fig. 3D). A complemented mutant stably expressing a copy of the *phoP* gene (Compl.) could restore intra-bacterial Asn level to that of WT-H37Rv. From these results, we conclude that *in vivo* Asn acquisition of *M. tuberculosis* requires the *phoP* locus. It should be noted that using infected THP-1 macrophages, we consistently noted a significantly lower intra-bacterial ammonia level for macrophage-derived *phoP*-KO strain relative to WT-H37Rv and a complemented strain (Fig. S3B). However, WT-H37Rv and *phoR*-KO, grown under identical *in vitro* conditions as described in Fig. 3C showed a comparable level of intra-bacterial ammonia (Fig. S3C).

With the uncovering of a role of *phoP* locus in mycobacterial Asn acquisition, we compared relative expression of representative Asn metabolism genes in *phoP*-KO mutant and WT-H37Rv, grown under low pH conditions (Fig. 3E). Our results demonstrate a significant down regulation of *ansP1* and *ansP2* expression in *phoP*-KO mutant relative to WT-H37Rv. Stable expression of a copy of *phoP* gene in the complemented mutant (Compl.) could rescue expression of *ansP1* and *ansP2* to the WT level. In contrast, expression of *ansA*, *asnT* and *glnA1* remain unaffected in *phoP*-KO mutant relative to WT bacilli. To assess whether transcription regulation of *ansP1* and *ansP2* is attributable to recruitment of PhoP, we also examined *in vivo* DNA binding of the regulator within the target promoters of WT-H37Rv, grown under normal and acidic conditions of growth. We utilized chromatin immunoprecipitation assays (ChIP) coupled with quantitative PCR measurements using anti-PhoP antibody (Fig. 3F). Our results highlight a significantly elevated recruitment of PhoP within the upstream regulatory regions of *ansP1* (ansP1up) and *ansP2* (ansP2up) for WT-H37Rv grown under acidic conditions relative to normal conditions.

These results are consistent with (a) importance of acidic pH-inducible activation of PhoP leading to high-affinity DNA binding/ transcription activation [28, 29], and (b) role of PhoP as a regulator of *ansP1* and *ansP2* expression. Table S1 enlists sequence of oligonucleotides used in RT-qPCR/ChIP-qPCR experiments. The region spanning upstream regulatory region of *aprA* (aprAup), which belongs to PhoP regulon, was used as a positive control and 16S rDNAup was used as an endogenous control.

### Direct transcriptional control of mycobacterial Asp and Asn transporters

With the results showing a considerable effect of PhoP on transcriptional control of *ansP1* and *ansP2*, we wished to investigate if PhoP functions as a direct regulator of *M. tuberculosis ansP1* and *ansP2* genes *in vivo.* To this end, we constructed transcriptional fusions to *lacZ* by cloning PCR-amplified fragments of *ansP1* and *ansP2* upstream regulatory regions at the ScaI site of pSM128, an integrative promoter probe vector for mycobacteria [30]. *M. smegmatis* harboring the transcriptional fusions, ansP1up-*lacZ* or ansP2up-*lacZ* (see methods for details) were co-transformed with pME1mL1-*phoP*, an inducible expression system expressing *M. tuberculosis* PhoP from the P_myc1_*tet*O promoter under TetR repressor as described previously [31]. Co-transformed cells were grown in 7H9 medium containing appropriate antibiotics with or without anhydrotetracycline (ATc; 50 ng/ml) to induce expression of PhoP. Notably, ansP1up-*lacZ* fusion was significantly activated with induction of PhoP at the 24-hour time point, as the β-galactosidase levels obtained in presence of ATc was ≍2.1±0.1**-**fold higher relative to the activity in the absence of ATc (Fig. 4A). Similarly, cells carrying ansP2up*-lacZ* fusion showed a ≍ 3.6±0.4-fold higher β-galactosidase activity when PhoP expression was induced compared to the uninduced sample (Fig. 4B). Note that for both cases in the absence of PhoP expression (pME1mL1 was used as an empty vector control), we failed to observe a significant difference in promoter activity in the absence or presence of ATc. In contrast, a strain harboring the reporter construct ansAup-*lacZ* (comprising –379 to +108 of the *ansA* upstream regulatory region relative to its ORF start site), under identical experimental conditions, failed to show promoter activation with induction of PhoP expression (1.0±0.02-fold change of β-galactosidase activity in induced versus uninduced samples; Fig. 4C). Insets compare expression of PhoP in *M. smegmatis* in the absence or presence of ATc. From these results, we conclude that *M. tuberculosis* PhoP directly activates *in vivo* expression of *ansP1* and *ansP2*. These results are consistent with the transcriptomic data suggesting PhoP-dependent activation of *ansP1* and *ansP2*, but not *ansA* expression (see Fig. 3E).

**Figure 4:**
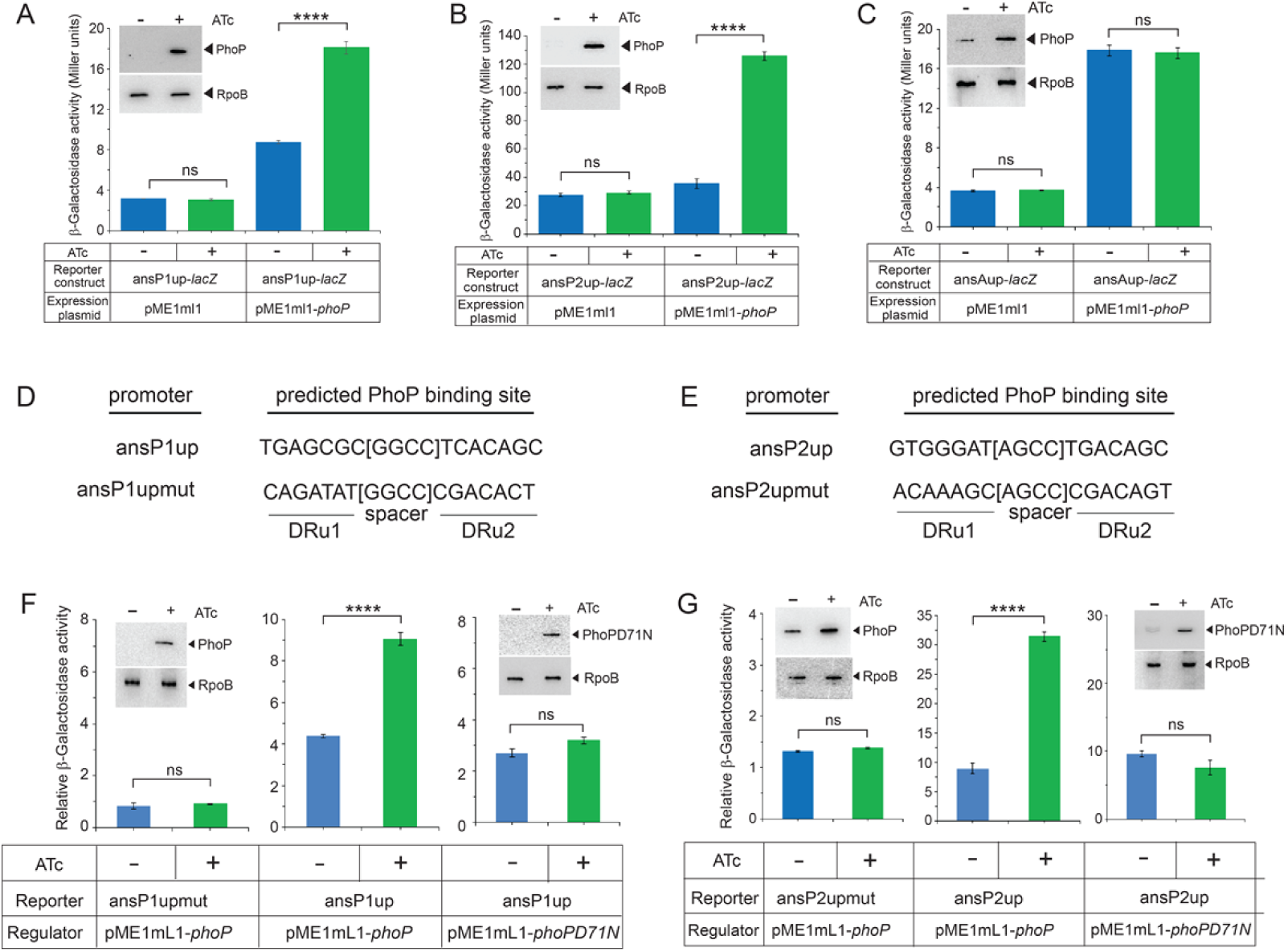
*M. tuberculosis* PhoP activates expression of *ansP1* and *ansP2 in vivo* in *M. smegmatis*. (A-C) *M. smegmatis* harbouring indicated fusion constructs along with PhoP expression plasmid or an empty vector were grown in the absence or presence of ATc and β-galactosidase activity from the transcription fusions were measured after 24 hours of growth. The values shown here are averages of at least three independent experiments with technical repeats (****P<0.0001; ns, non-significant difference). Insets compare expression of PhoP in crude extracts containing equal amounts of total protein (as determined by Bradford assay) by Western blot analysis using anti-PhoP antibody (Bio-Bharati). As a loading control, identical extracts were probed with antibody against β-subunit of RNA polymerase (RpoB, Abcam). (D-E) These panels show newly identified PhoP binding direct-repeat motifs within ansP1up and ansP2up, respectively. To examine the importance of PhoP binding motifs within the *ansP1* and *ansP2* promoters, mutations in the PhoP binding sites were introduced as shown in the figure (ansP1upmut and ansP2upmut, respectively). (F-G) The reporter assays were carried out using indicated wild-type and mutant promoters along with inducible expression of WT/mutant PhoP proteins and data analysed as described in the methods. Note that phosphorylation-defective PhoPD71N was used to examine the effect of phosphorylation of PhoP on transcription activation of promoters. The corresponding insets compare expression of the mutant PhoP protein in comparable amounts of crude lysates with RpoB as the loading control.

To identify the mechanism of PhoP-dependent activation, we next probed PhoP binding sites within the target promoters (ansP1up, and ansP2up) using previously reported consensus PhoP binding site. Figs. 4D-E show predicted PhoP binding sites. To investigate role of the newly-identified sites in PhoP-coupled transcription activation of *ansP1* and *ansP2*, changes were introduced in the nucleotide sequence of the upstream and downstream repeat units of ansP1up and ansP2up by replacing purines with non-pairing pyrimidines (and *vice versa*) and by inverting the downstream repeat units as shown in the figures. The promoter fragments carrying changes in the PhoP binding sites (ansP1upmut and ansP2upmut, respectively) were subsequently cloned in pSM128 and used as transcriptional fusions to examine regulatory effect of PhoP. Interestingly, with induction of PhoP expression we observed no significant changes in the level of relative β-galactosidase activities from the promoter(s) carrying changes at the PhoP binding sites. Our results show that the fold change of β-galactosidase activity at 24 hours in the presence and absence of inducing PhoP expression was 1. 1(±0.12)-and 1.05(±0.02)-fold (compare green versus blue columns) from the ansP1upmut-*lacZ* and ansP2upmut-*lacZ*, respectively (left panels of Figs. 4F-G). The insets compare expression of PhoP in the absence and presence of ATc.

These results are in striking contrast with PhoP-dependent 2.1(±0.1)-and 3.6(±0.4)-fold activation of ansP1up and ansP2up expression, respectively, under identical conditions examined (middle panels, Figs. 4F-G). Thus, we conclude that PhoP binding at the newly-identified sequence motif within the corresponding regulatory regions is necessary and sufficient for transcription activation of *ansP1*, and *ansP2*, respectively. To examine the effect of phosphorylation of PhoP on activation of ansP1up and ansP2up, we utilized identical fusion constructs along with inducing expression of PhoPD71N, a mutant PhoP defective for phosphorylation at Asp-71 [29]. Although PhoPD71N displayed an ATc-dependent induction of expression comparable to that of wild-type PhoP (insets to right panels, Figs. 4F-G), it showed 1.2±0.07 and 0.8±0.14-fold variation of promoter activation from ansP1up, and ansP2up, respectively (right panels, Figs. 4F-G). Thus, we conclude that PhoP-dependent activation of *ansP1*, and *ansP2* is attributable to phosphorylation of PhoP. These results are consistent with low pH-dependent elevated DNA binding of P∼PhoP during mycobacterial growth under acidic conditions (relative to normal conditions), known to activate functioning of the regulator via phosphorylation [23, 29].

### PhoP-dependent Asn utilization is essential for intracellular survival of mycobacteria

To examine the effect of over-expression of *ansP1* and *ansP2* in the *phoP*-KO background, the ORFs were cloned within the multicloning sites of mycobacterial expression vector p19KPro (see methods) [32] and fold change in expression under normal conditions was verified by determining the mRNA level (Figs. S4A-B). *phoP*-KO::*ansP1* and *phoP*-KO::*ansP2*, over-expressing *ansP1* and *ansP2*, respectively, were grown under acidic conditions in the presence of Asn as the sole source of nitrogen. Upon CFU enumeration, we observed that at day ‘0’ both over-expressing constructs grew comparably well without any significant difference with *phoP*-KO harboring the empty vector (p19KPro) (Fig. 5A). However, at day 7, we observed a significantly improved growth of both the over-expressing strains compared to *phoP*-KO mutant.

**Figure 5:**
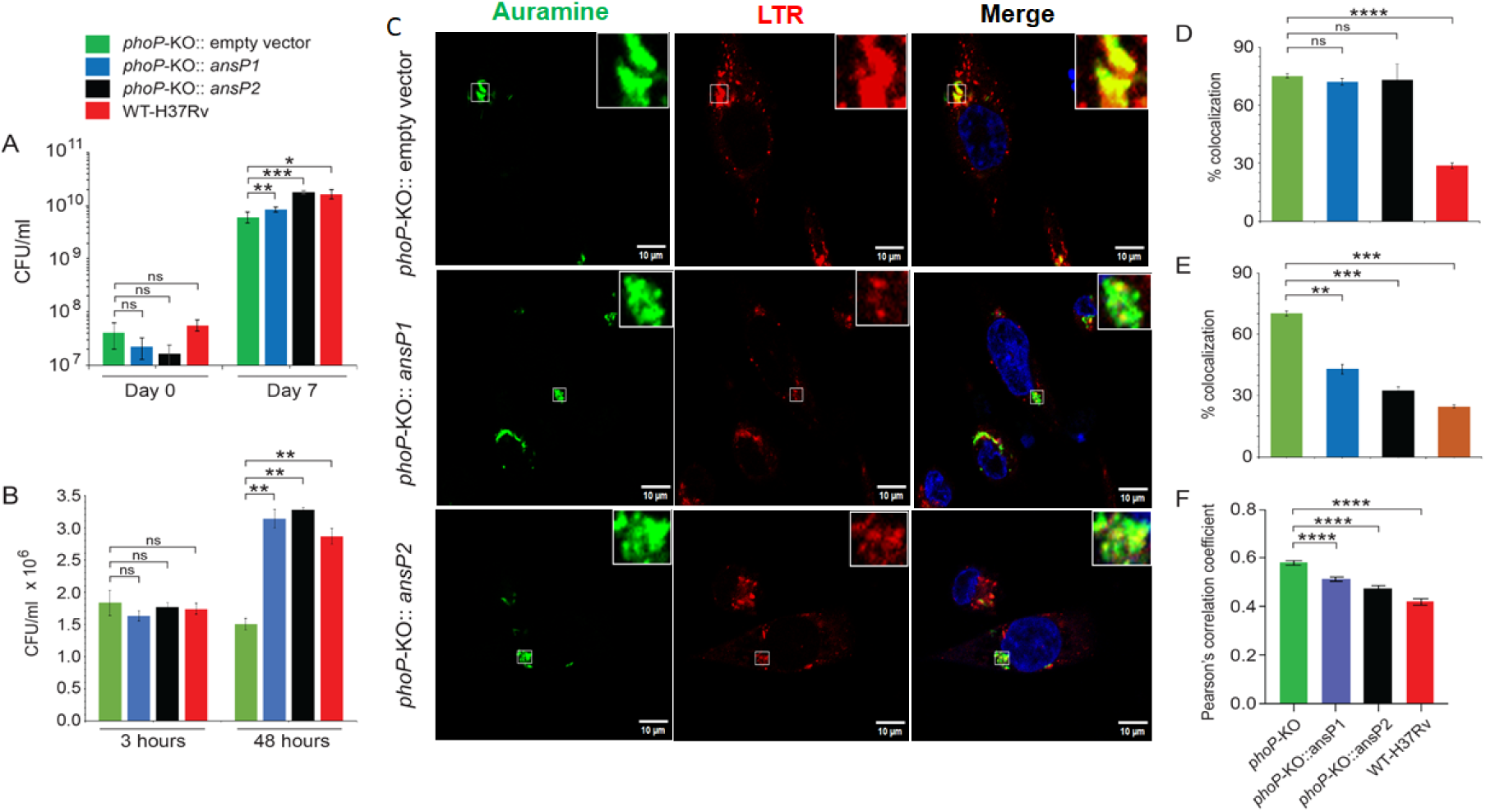
*ansP1* and *ansP2* over-expressions promote intracellular survival of *M. tuberculosis*. *phoP*-KO was transformed with *ansP1* and *ansP2* over-expressing plasmids to construct the over-expressing strains as described in the methods. (A) To compare their relative abilities to utilize Asn, these strains were grown *in vitro* under acidic conditions (pH 4.5) and in the presence of Asn as the sole nitrogen source and CFU values enumerated. (B) To examine mycobacterial survival in cellular models, THP-1 macrophages were infected with indicated mycobacterial strains, and 3-hour and 48-hour post infection intracellular bacterial CFU were enumerated. (C) To examine effect of over-expression of genes on bacterial colocalization, THP-1 macrophages were infected with indicated bacterial strains. Mycobacteria and host cells were stained with phenolic auramine solution, and LysoTracker (LTR) respectively. Host cell nuclei were made visible by Hoechst dye. Two fluorescence signals (mycobacterial strains: green; and lysosomes: red) and their merge with host nuclei in blue are displayed by confocal images (scale bar: 10 µm). (D-E) These panels show CFU enumeration of intracellular bacteria for (D) 3-hour and (E) 48-hour post infection samples, respectively. (F) In case of 48-hour post infection samples, the data represent Pearson’s correlation coefficient of images displaying internalized auramine-labelled mycobacteria and Lysotracker red marker in macrophages and were evaluated using image-processing software NIS elements (Nikon). The results represent average data from multiple biological replicates (****P<0.0001; ***P<0.001; **P<0.01; *P<0.05; ns, non-significant difference).

In fact, growth of *phoP*-KO::*ansP2* was almost comparable to WT-H37Rv. Likewise, both *phoP*-KO::*ansP1* and *phoP*-KO::*ansP2* displayed significantly improved bacterial growth within THP-1 macrophages compared to the *phoP*-KO mutant (Fig. 5B), suggesting that independent over-expression of *ansP1* and *ansP2* in *phoP*-KO mutant restored mycobacterial stress tolerance both *in vitro* and *in vivo*.

We next studied phagosome-lysosome fusion in THP-1 macrophages using these mycobacterial strains (Fig. 5C). In this assay, mycobacteria and host cells were stained with phenolic auramine solution, and Lysotracker (LTR), respectively. Host cells nuclei were made visible by Hoechst dye and the two fluorescent signals and their merge with host nuclei in blue are shown by confocal images (scale bar: 10 µm). Consistent with previous studies [33], WT bacilli could inhibit phagosome maturation (Fig. S4C), whereas infection of macrophages with *phoP*-KO harboring the empty vector readily matured into phagolysosomes (top panel, Fig. 5C), suggesting increased trafficking of the mutant bacilli to lysosomes. Importantly, 48-hour post-infection, both the *phoP*-KO::*ansP1* (middle panel, Fig. 5C) and *phoP*-KO:: *ansP2* (bottom panel, Fig. 5C) independently showed significantly reduced co-localization with lysotracker red (LTR) compared to *phoP*-KO mutant, suggesting reduced phagosome maturation by *phoP*-KO::*ansP1* and *phoP*-KO::*ansP2* relative to *phoP*-KO. Note that under identical conditions for 3-hour post-infection, both the over-expressing constructs showed comparable phagosome maturation as that of *phoP*-KO mutant (Fig. 5D). Fig. 5E shows percent co-localization of mycobacterial strains with LTR, 48-hour post infection. The Pearson’s correlation co-efficient values, as quantified from co-localization signals, are plotted in Fig. 5F. These results are consistent with improved bacterial survival within macrophages by the over-expressing constructs compared to *phoP*-KO (see Fig. 5B). Together, we conclude that ectopic expression of *ansP1* and *ansP2* contribute to inhibition of phagosome maturation, arguing in support of a functional role of PhoP in regulating mycobacterial phagosome-lysosome fusion.

## Discussion

Identifying mycobacterial acquisition of essential elements hold a key step toward understanding of host-pathogen interactions in TB. Previous studies suggest that Asp and Asn are key nitrogen sources used by *M. tuberculosis* during infection [16, 17, 20]. Further, it has been shown that Asn is an effective nitrogen source of the pathogen via transport through the amino acid permease AnsP2, and subsequent hydrolysis by AnsA, the asparaginase. These results are of significant physiological relevance given the fact that human plasma contains 50-60 µM Asn [34], and its concentration is much higher in white blood cells [35, 36]. In addition, Asn has been shown to accumulate in mycobacterial vacuoles inside infected macrophages [20], suggesting *in vivo* abundance of the amino acid to the intracellular pathogen during infection.

However, the regulatory network of the pathogen that mediates effective utilization of Asn remains unknown.

The fact that Asn, unlike Gln or ammonium, effectively supports *in vitro* growth of WT bacilli under low pH conditions of growth (Figs. 1 and S1), is remarkably consistent with previous work by Nyrolles and coworkers showing Asn-dependent mycobacterial resistance to acidic pH [20]. These results connect Asn acquisition with the virulence regulator PhoP, a major player in phagosomal acidic pH dependent adaptive program of mycobacteria [21, 22, 37].

Subsequent transcriptomic studies uncover PhoP-dependent regulation of Asp transporter (AnsP1) and Asn transporter (AnsP2), respectively (Fig. 3). These results unambiguously suggest functioning of the regulator as a modulator of Asn acquisition. To probe the *in vivo* significance, we compared intra-bacterial pH of WT-H37Rv and a mutant lacking *phoP*, grown under acidic pH (pH 5.5) and in the presence of Asn as the sole nitrogen source. Interestingly, *phoP*-KO mutant unlike WT-H37Rv failed to maintain intra-bacterial pH (Fig. 3A) and displayed a significantly reduced intra-bacterial Asn level (Fig. 3B). These results are consistent with PhoP-dependent Asn uptake contributing to mycobacterial pH homeostasis. PhoP-controlled activation of *ansP1* and *ansP2* followed a direct regulatory mechanism as shown by specific *in vivo* recruitment of the regulator within the upstream regulatory regions of *ansP1* and *ansP2* (Fig. 2C) coupled with reporter assays showing promoter activation in *M. smegmatis* (Fig. 4). Further, to assess the effect of Asn acquisition on mycobacterial growth, our subsequent experiments uncover that *ansP1* and *ansP2* over-expression in a *phoP*-KO background significantly influenced mycobacterial growth both *in vitro* and within host cells (Fig. 5).

Together, we conclude that growth attenuation of *phoP*-KO in host cells is attributable, at least in part, to the mutant’s failure to activate *ansP1* and *ansP2* transporters for efficient utilization of Asn, leading to reduced stress tolerance under phagosomal acidic conditions. These findings have significant implications on the mechanism of mycobacterial adaptive response at the host pathogen crossroads.

A previous study showed that *ansP2*-KO mutant was partially impaired in nitrogen incorporation from Asn *in vitro* and was not attenuated *in vivo* [20]. Thus, AnsP1, an AsnP2 paralogue was considered to complement AnsP2 functionality [38]. However, *ansP1*-KO grew comparably well as that of WT-H37Rv with Asn as the sole nitrogen source and transported Asn *in vitro* as effectively as that of WT-H37Rv[16]. These results suggest that *ansP1* and *ansP2* appear to complement each other, at least in part, and only obliteration of both genes simultaneously would result a clean phenotype. These results explain and fit well with (a) over-expression of either *ansP1* or *ansP2* in a *phoP*-KO facilitating mycobacterial utilization of Asn both *in vitro* and within host cells, and (b) growth attenuation of *phoP*-KO, lacking the key regulator which activates expression of both *ansP1* and *ansP2*. It should be noted that functioning of PhoP is critically required for *M. tuberculosis* virulence [39, 40] as it impacts numerous aspects of bacterial physiology including complex lipid biosynthesis [41–43], early response to hypoxia [44], regulation of ESAT-6 secretion [44–47], and pH sensing during intracellular adaptation [21–23]. Using genome-wide probing, SELEX studies and ChIP-seq analysis clearly identified the consensus PhoP binding site [33, 48]. Results reported here showing mutagenesis of promoters coupled with *ex vivo* reporter assays using transcriptional fusion of WT and mutant promoters (carrying changes at the PhoP binding sites) establish that the newly-identified direct repeat motif alone is likely attributable to PhoP-DNA interactions at the regulatory sites, and recruitment of P∼PhoP at these sites is essential for PhoP-dependent regulation of *ansP1*, and *ansP2* (Fig. 4). Consistent with these results, the PhoP-DNA co-crystal structure demonstrates tandem head-to-head binding of the proteins to the adjacent repeat units where a single protomer recognizes each unit [49–51]. Interestingly, our results on newly-identified PhoP binding sites within the *ansP1* and *ansP2* promoters are in broad agreement with the presence of two 7-bp repeat units separated by a 4-bp spacer sequence (Figs. 4D-E).

However, the nucleotide sequences of the repeat sites are significantly different from each other, suggesting how similar sequence modules with variable nucleotide sequences, display differential affinity of the same transcription factor (PhoP), and account for functional diversity to accommodate variable physiological functions.

The above results expand upon our previous work on sequence specific promoter recognition by P∼PhoP to demonstrate transcriptional control of *ansP1* and *ansP2*, genes with related metabolic function involving Asp and Asn acquisition of mycobacteria. In agreement with *in vivo* regulation by P∼PhoP (Fig. 3E), our results show a significantly improved *in vivo* DNA binding of PhoP under acidic conditions of mycobacterial growth (Fig. 3F). These results fit well with and explain acidic pH-dependent activation of PhoP via PhoR-dependent sensing [23]. In keeping with this, reporter assays fail to show promoter activation by PhoPD71N (Figs. 4F-G, right panels), a phosphorylation-defective mutant protein used to circumvent the problem of PhoP being phosphorylated *in vivo* by a non-cognate sensor kinase. Together, these results as shown schematically (Fig. 6), uncover that (a) direct interactions between the P∼PhoP and the newly-identified PhoP binding site(s) are essential for activation of target genes *ansP1* and *ansP2*, and (b) phosphorylation of a single residue of PhoP contributes to regulation of Asn acquisition and maintenance of pH homeostasis, a novel result of unusual significance at the host-pathogen cross-roads. In conclusion, these results provide yet another example of close connectivity shaped through evolution to accommodate physiology and virulence regulation of microbial pathogens together. It also underscores the need to further extend exploration of such connections and identify novel strategies to combat infectious diseases.

**Figure 6:**
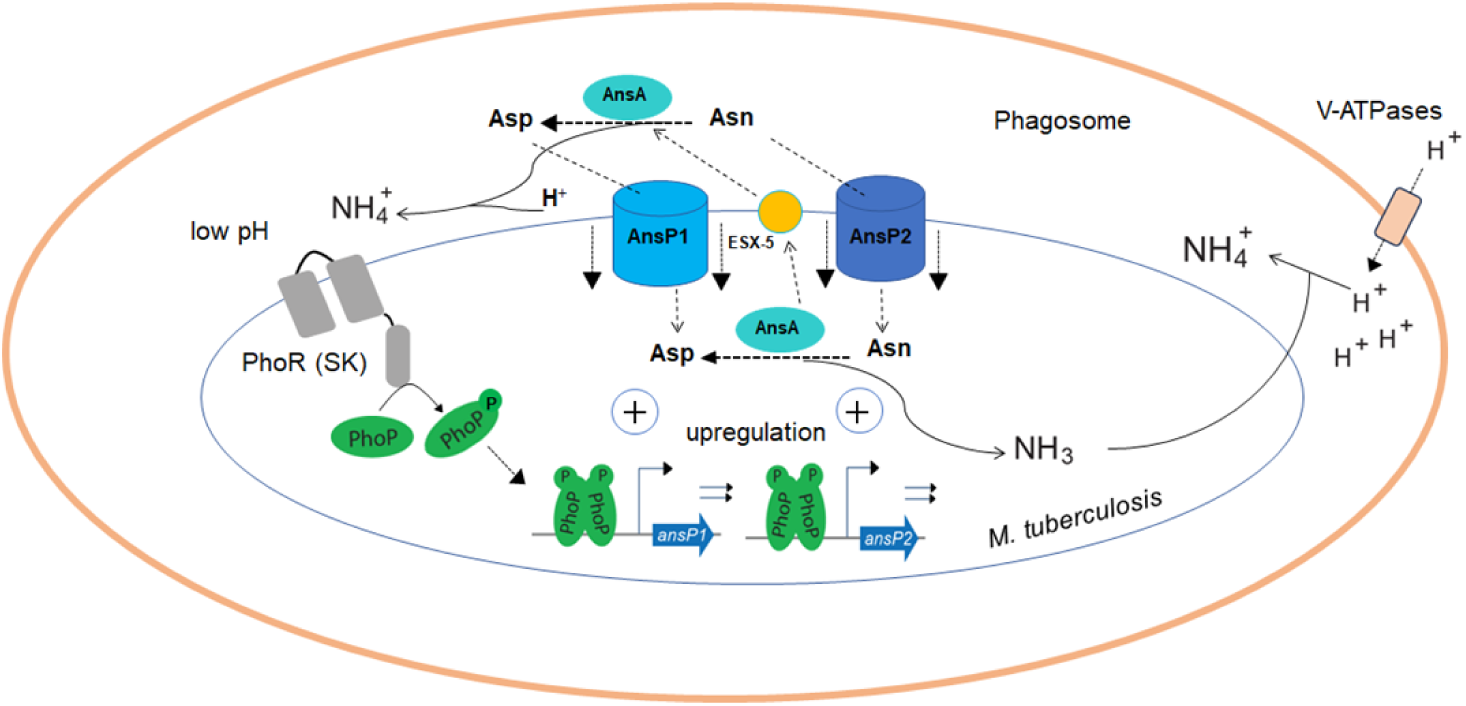
Model showing a schematic representation of how Asp/Asn acquisition couple mycobacterial tolerance to acidic pH. Within macrophages Asp and Asn are captured by *M. tuberculosis* AsnP1 and AnsP2, respectively. Cytosolic AnsA hydrolyzes Asn to Asp with release of ammonia. While Asn is transported by AnsP2 and contributes to mycobacterial nitrogen assimilation, released ammonia enables phagosomal pH buffering [17]. Importantly, transcriptional activation of *ansP1* and *ansP2* under acidic pH is attributable to low pH dependent PhoR-mediated activation of PhoP, followed by PhoP functioning as an activator by direct recruitment of P∼PhoP within the upstream regulatory regions of *anP1* and *ansP2*, respectively. Together, these events effectively integrate nitrogen metabolism and pH homeostasis to help survive mycobacteria in a hostile intracellular environment.

## Materials and Methods

### Bacterial strains and culture conditions

*E. coli* TOP10 and DH5α strains, used for cloning, were grown on LB agar medium and/or LB broth. *M. tuberculosis* H37Rv and *M. smegmatis* mc^2^155 were cultured in Middlebrook 7H9 broth supplemented with 10% OADC (oleic acid, albumin, dextrose, and catalase) and 0.05% Tween-80 or on Middlebrook 7H11 medium supplemented with 10% OADC. Additional details on mutant strains such as *phoP*-KO, the complemented mutant, and the knock-down mutants (*phoP*-KD and *prrB*-KD) are available in earlier studies [23, 26, 41]. When required, mycobacterial strains were transformed and selected in the presence of specific antibiotics at the following concentrations: hygromycin (50 µg/ml), kanamycin (20 µg/ml) and streptomycin (20 µg/ml).

For nitrogen-defined conditions, synthetic m7H9 medium lacking arginine and glutamic acid was used, ammonium sulfate and ferric ammonium citrate replaced with sodium sulfate and ferric citrate, respectively, and mycobacterial cells grown in the presence of specific amino acids as the sole nitrogen sources. Colony-forming units (CFUs) were enumerated by inoculating the strains into fresh 7H9-ADS (albumin-dextrose-sodium chloride) medium at an OD_600_ of ∼0.05. Cultures at the onset (0 day) and after 7 days of growth were serially diluted, and plated at various dilutions on 7H11-OADS agar plates for 21 days to enumerate CFU. For mycobacterial growth under specific stress conditions, strains were grown to mid-log phase (OD_600_ 0.4 to 0.6) and exposed to different stress conditions as described previously [52]. For *in vitro* growth curve experiments, strains were diluted to an initial OD_600_ of 0.05 in Middlebrook 7H9 medium, and the OD_600_ was recorded at different time points over a period of 10 days. In each case, data from three independent experiments were used to generate the growth curve.

### Construction of *M. tuberculosis* knock-down mutants

We used a previously reported CRISPRi system [25] to construct knock-down mutants (*phoP*-KD and *prrB*-KD), as described previously [23, 26]. This method effectively inhibits expression of target genes via inducible expression of dCas9 along with gene specific guide RNAs (sgRNA). To compare expression of target genes, the mutant strains were verified by RT-qPCR using total RNA from the respective knock-down constructs relative to corresponding uninduced strains.

### Determination of intra-bacterial pH and Asn level

Indicated mycobacterial strains were transformed with pH-GFP, a plasmid encoded pH-sensitive green fluorescent protein (pH-GFP) as a pH probe [27], and grown in synthetic m7H9 supplemented with Asn or ammonium as the sole source of nitrogen as described above. Cultures in early log phase were adjusted to an OD_600_ of 0.5, 1 ml aliquots in triplicates were further incubated at 37°C and concentrated 5-fold to enhance GFP signal intensity. Next, 100 µl of cell aliquots were used in a fluorescent plate reader (Molecular Device) with excitation wavelengths set at 395 nm and 475nm, and recording emission at 510 nm, respectively. pH values were determined through absorbance ratios (395 nm/475nm) against a standard curve as described previously [27].

Intra-bacterial Asn levels were measured by growing WT and *phoP*-KO mutant to an OD_600_ of 0.1. After reaching this OD, the cultures were incubated at 37°C for an additional three days, and a comparable number of WT-H37Rv and mutant cells lysed in a buffer containing 50 mM Tris (pH 7.5), 1% Triton X-100, 150 mM NaCl, 1 mM PMSF, 1 mM EDTA and 5% glycerol, and Lysing Matrix B (100 µm silica beads) in a bead beater (MPBio) at a speed setting of 6.0 for 1 minute, with cooling intervals between cycles to prevent overheating. After repeating the procedure for 3 cycles, the lysates were clarified at 1300 rpm for 10 minutes, and supernatants collected. 20 µl of each sample was taken into wells of a 96-well flat-bottom plate in duplicates, and intra-bacterial Asn levels were determined by using the Megazyme reagents and protocol according to the manufacturer’s recommendation.

### Bacterial RNA extraction

Total RNA was isolated from bacterial cultures grown under specific conditions as described earlier [23, 26]. In brief, 25 ml of bacterial culture (OD_600_ = 0.6-0.8) was mixed with equal volume of a 5 M guanidinium thiocyanate solution containing 1% β-mercaptoethanol and 0.5% Tween-80. The cells were pelleted by centrifugation, resuspended in 1 ml of Trizol (Ambion), and then subjected to lysis with Lysing Matrix B (100 µm silica beads, MPBio). Lysis was performed using a FastPrep-24 bead beater (MP Bio) at a speed setting of 6.0 for 30 seconds, repeating the process for three cycles with cooling intervals to prevent overheating. The resulting lysates were centrifuged at 13,000 rpm for 10 minutes, and the RNA was extracted from the supernatant using the Direct-Zol™ RNA isolation kit (ZYMO). Genomic DNA contamination was removed by treatment with DNase I (Promega). RNA quality was assessed using an UV-VIS spectrophotometer (Shimadzu Corporation), and the integrity of 23S and 16S rRNA was further confirmed by gel electrophoresis. RNA concentrations were determined with a Qubit fluorometer (Invitrogen).

### Quantitative Real-Time PCR Analysis

Transcriptional profiling was performed using RNA extracted from *M. tuberculosis* cultures grown under specific growth conditions. cDNA synthesis and PCR amplification were performed with the Superscript III Platinum SYBR Green One-Step RT-qPCR Kit (Invitrogen), following previously reported protocol [52]. To determine fold changes in gene expression, multiple independent RNA preparations were analyzed by ΔΔCT method [53] using *M. tuberculosis rpoB* or *16S rDNA* as internal controls. The primer sequences used for the RT-qPCR analysis are provided in Table S1.

### Cloning

To over-express mycobacterial *ansP1* and *ansA*, we amplified both the ORFs from *M. tuberculosis* genomic DNA using primer pairs FPansP1/RPansP1 and FPansA/RPansA, respectively, and cloned between BamHI and HindIII sites of the mycobacterial expression vector p19Kpro [32]. Similarly, *ansP2* was amplified by using FPansP1/RPansP2 primer pair and cloned between HindIII and ClaI sites of p19Kpro. Recombinant His_6_-tagged PhoP was cloned between NdeI and HindIII sites of pET-28b using the primer pair phoPstart/phoPstop. To express *M. tuberculosis* PhoP in mycobacteria, the ORF with an N-terminal Flag-tag was cloned, and expressed from p19Kpro [32] as described previously [47]. To express *M. tuberculosis* PhoP in *M. smegmatis*, wild-type or mutant *phoP* was cloned in pME1mL1 [31] and expressed from P_myc1_*tet*O promoter under TetR repressor, as described elsewhere [29]. Each construct was checked by DNA sequencing. Tables S2 and S3 list the oligonucleotide primers and plasmids, respectively, used for cloning reported in this work.

### Macrophage Infection Assays

Infection of THP-1 macrophages with *M. tuberculosis* strains was carried out at a multiplicity of infection (MOI) of 1:10, according to previously established protocols [33]. *M. tuberculosis* H37Rv strains were stained using phenolic auramine solution, while macrophage cells were labeled with Lyso-Tracker Red (Invitrogen; 150 nM). Following infection, cells were fixed and examined using a Nikon A1R confocal microscope. Laser and detector settings were optimized based on macrophages infected with WT-H37Rv to ensure consistency, and images processed digitally with IMARIS software (version 9.20), with standard intensity thresholds. The percentage of bacterial co-localization was calculated by analyzing 50 infected macrophages from 10 different fields across independent replicated experiments.

To determine intra-bacterial Asn level of macrophage-derived *M. tuberculosis* strains, THP-1 macrophages were infected with WT-H37Rv and *phoP*-KO mutant at an MOI of 1:10. 48-hour post infection, macrophages were lysed with 0.06% SDS in 1X PBS and centrifuged at 4000 rpm for 10 minutes. The pellet was washed twice with 1X PBS buffer to ensure complete removal of macrophage debris. The bacterial pellet was then resuspended in 0.2 ml of 1X PBS and comparable number of bacterial cells from each strain was lysed with IP buffer (50 mM Tris (pH 7.5), 150 mM NaCl, 1 mM EDTA, 1% Triton X-100, 1 mM PMSF and 5% glycerol) containing protease inhibitor cocktail. The resultant lysates were centrifuged at 13000 rom for 10 minutes at 4°C. The supernatants were then used to measure intra-bacterial Asn level as described above.

### Chromatin Immunoprecipitation (ChIP)-qPCR

ChIP assays were performed with actively growing *M. tuberculosis* cultures as described previously [23, 52] using anti-FLAG/ anti-PhoP antibody and protein A/G agarose beads (Pierce). For qPCR measurements, immunoprecipitated DNA was diluted by 10-fold and combined with a reaction mixture with SYBR Green mix, appropriate primers (Table S1) targeting specific promoter regions, and Platinum Taq DNA polymerase (1 Unit, Invitrogen). Amplification was done using a real-time PCR detection system (Applied Biosystems). Control primers specific to *16S rDNA* or *rpoB* genes were included to validate specific enrichment from IP samples. To assess recruitment efficiency, mock IP samples (without antibody) were used as negative controls, and data were normalized to PCR signal from the control reactions. Melting curve analysis confirmed amplification of a single product in all reactions. Replicate qPCR measurements were performed using DNA from multiple independent bacterial cultures. PhoP binding to target promoters was evaluated by analyzing 1 µl of IP or mock IP DNA, combined with SYBR green mix (Invitrogen) and promoter-specific primers.

### Construction of *lacZ* fusions

To construct the promoter-*lacZ* fusions, upstream of *ansP1* (encompassing-370 to +95 of *ansP1*), *ansP2* (encompassing-384 to +88 of *ansP2*), and *ansA* (encompassing-379 to +108 of *ansA*, relative to corresponding ORF start sites were PCR amplified using WT-H37Rv genomic DNA as template and FPansP1up/RPansP1up, FPansP2up/RPansP2up and FPansAup/RPansAup, as primer pairs, respectively (Table S2). The PCR-amplified fragments were cloned upstream of *lacZ* in the promoter-less integrative plasmid pSM128 [54], a kind gift from Prof. Issar Smith, Public Health Research Institute, UMDNJ. These plasmids were introduced in *M. smegmatis* by electroporation and co-transformants with pME1mL1/pME1mL1-*phoP* were selected on 7H10 plates containing hygromycin (hyg) and streptomycin (strep). To assess the role of PhoP binding sites, sequences of the repeat units within ansP1up, and ansP2up were mutated using overlap extension method by interchanging the As with Cs, the Gs with Ts, and *vice versa* as shown in Figs. 4D-E. All the *lacZ* transcriptional fusions were verified by DNA sequencing.

### Regulation of mycobacterial promoter activity in *M. smegmatis*

Inducible expression of *M. tuberculosis* PhoP (WT/mutant) in *M. smegmatis* from pME1mL1, and measurement of WT/mutant promoter activities from co-transformed strains, as indicated in the figure, were carried out as described previously [29]. A_420_ values of the supernatant were determined, and units of β-galactosidase calculated as follows: 1000 x A_420_ per mg protein per minute in Miller Units. The protein concentration in the cell lysates were measured, and *M. smegmatis* co-transformed with pSM128 and pME1ml1 served as the control. PhoP expression from crude lysates of *M. smegmatis* were detected by Western blot analysis using anti-PhoP antibody.

### Statistical analysis

Data are presented as arithmetic means of the results obtained from multiple replicate experiments ± standard deviations. Statistical significance was determined by Student’s paired t-test using Microsoft Excel or Graph Pad Prism. Statistical significance was considered at P values of 0.05 (*P≤0.05; **P≤0.01; ***P≤0.001; ****P≤0.0001).

## Data availability statement

The data reported in this manuscript are part of the main text and its Supporting Information files.

## Acknowledgements

This study was supported by funding from CSIR-IMTECH intramural grant OLP-0170, CSIR-funded project FBR070308, and SERB-funded project (CRG/2023/00109) to D.S. B. B., and K.M. were supported by CSIR, and Kajal was supported by DST Inspire fellowship. The funders had no role in study design, data collection and analysis, decision to publish or preparation of the manuscript.

We acknowledge G. Marcela Rodriguez and Issar Smith (PHRI, New Jersey Medical School - UMDNJ) for *phoP*-KO mutant, the complemented mutant strain, plasmid pSM128, and Sabine Ehrt (Weil Medical College of Cornell University) for pME1mL1 expression vector.

## Author Contribution

**Conceptualization:** Bhanwar Bamniya, Khushboo Mehta, and Dibyendu Sarkar

**Data curation:** Bhanwar Bamniya, Kajal, Khushboo Mehta

**Formal analysis:** Bhanwar Bamniya, Kajal, Khushboo Mehta, Dibyendu Sarkar

**Funding acquisition:** Dibyendu Sarkar

**Investigation:** Bhanwar Bamniya, Kajal, Khushboo Mehta, Dibyendu Sarkar

**Methodology:** Bhanwar Bamniya, Kajal, Khushboo Mehta

**Project administration:** Dibyendu Sarkar

**Resources:** Bhanwar Bamniya, Kajal, Khushboo Mehta, Dibyendu Sarkar

**Software:** Bhanwar Bamniya, Kajal, Khushboo Mehta, Dibyendu Sarkar

**Supervision:** Bhanwar Bamniya, Dibyendu Sarkar

**Visualization:** Bhanwar Bamniya, Kajal, Khushboo Mehta, Dibyendu Sarkar

**Writing – original draft:** Dibyendu Sarkar

**Writing – review & editing:** Bhanwar Bamniya, Dibyendu Sarkar

## Supporting Information

**Figure S1:** To examine mycobacterial utilization of (A) ammonium salt or (B) Asp as the sole nitrogen source, WT-H37Rv was grown in synthetic m7H9 medium supplemented with indicated nitrogen source at pH 6.6 and pH 4.5, respectively. Panels C and D represent the enumerated CFU values of corresponding growth experiments at day 0 and day 7, respectively (*P≤0.05; **P≤0. 01; ns, nonsignificant difference). The values represent average of three independent growth experiments.

**Figure S2:** (A) To investigate the role of *phoP* locus in mycobacterial utilization of ammonium salt, WT-H37Rv and the indicated knock-down mutants were grown in synthetic m7H9 media supplemented with ammonium salt as the sole nitrogen source under normal pH. (B) Relative mRNA levels were measured in *prrB*-KD mutant grown under normal conditions using RT-qPCR. *prrB* expression level was significantly reduced in the knock-down mutant (**P< 0.01), whereas *phoP* expression of the mutant was comparable to that of WT-H37Rv (ns, nonsignificant difference); 16S rDNA was used as an endogenous control. (C) To further verify the role of *phoP* locus in mycobacterial utilization of ammonium salt, WT-H37Rv, *phoP*-KO mutant and complemented strain were grown under normal conditions in synthetic m7H9 media supplemented with ammonium salt as the sole nitrogen source. In all cases, growth experiments represent average values from biological triplicates.

**Figure S3:** (A) WT-H37Rv and *phoP*-KO mutant were grown *in vitro* under acidic pH conditions (pH 4.5) using synthetic m7H9 medium supplemented with Asn as the sole source of nitrogen, and intra-bacterial ammonia levels were measured as described in the methods. (B) In this experiment, THP-1 macrophages were infected with indicated strains of *M. tuberculosis*.

Next, 48-hour post-infection bacterial strains were isolated from infected macrophages, and intra-bacterial ammonia levels measured as described in the methods. The experiments were performed in biological triplicates and average values plotted with standard deviation (***P<0.001; **P<0.01; *P<0.05; ns, non-significant difference).

**Figure S4:** Over-expression of PhoP-controlled (A) *ansP1*, and (B) *ansP2* in the *phoP*-KO mutant. Expression levels of indicated genes were determined using corresponding over-expression strains, constructed by cloning indicated ORFs (see methods for further details). Fold change in expression was evaluated by RT-qPCR relative to 16S rDNA level of *phoP*-KO mutant harbouring the empty vector (normalized to 1.0 and represented by empty columns). The results show average values from biological triplicates, each with two technical repeats (**P≤0.01; ***P≤0.001). Nonsignificant differences are not shown. (C) THP-1 macrophages were infected with WT-H37Rv as described in the methods. Mycobacteria and host cells were stained with phenolic auramine solution, and LysoTracker respectively. Host cell nuclei were made visible by Hoechst dye. Two fluorescence signals (mycobacterial strains: green; and lysosomes: red) and their merge with host nuclei in blue are displayed by confocal images (scale bar: 10 µm).

**Table S1:** Sequence of oligonucleotide primers used for RT-qPCR and ChIP-qPCR experiments reported in this study

**Table S2:** Oligonucleotide primers used for cloning and amplifications reported in this study

**Table S3:** Plasmids used for cloning and amplifications reported in this study

